# GelMap: Intrinsic calibration and deformation mapping for expansion microscopy

**DOI:** 10.1101/2022.12.21.521394

**Authors:** Hugo G.J. Damstra, Josiah B. Passmore, Albert K. Serweta, Ioannis Koutlas, Mithila Burute, Frank J. Meye, Anna Akhmanova, Lukas C. Kapitein

## Abstract

Expansion microscopy (ExM) is a powerful technique to overcome the diffraction limit of light microscopy by physically expanding biological specimen in three dimensions. Nonetheless, using ExM for quantitative or diagnostic applications requires robust quality control methods to precisely determine expansion factors and to map deformations due to anisotropic expansion. Here we present GelMap, a flexible workflow to introduce a fluorescent grid into pre-expanded hydrogels that scales with expansion and reports deformations. We demonstrate that GelMap can be used to precisely determine the local expansion factor and to correct for deformations without the use of cellular reference structures or pre-expansion ground truth images. Moreover, we show that GelMap aids sample navigation for correlative uses of expansion microscopy. Finally, we show that GelMap is compatible with expansion of tissue and can be readily implemented as a quality control step into existing ExM workflows.

## INTRODUCTION

In Expansion Microscopy (ExM) the effective resolution of light microscopy is increased by physical expansion of cells and tissues (Chen et al., 2015). Biological specimens are anchored to a swellable hydrogel, typically by conjugation of an acrylate group to free lysines and subsequent polymerization of an acrylamide gel that includes sodium acrylate. Next, the sample is chemically homogenized to detach the gel from the culturing substrate and prevent resistance during expansion. Finally, addition of water induces swelling in all dimensions (Tillberg et al., 2016). Since ExM can be used to directly expand cells and intact tissues, ExM has quickly become an important technique in biological research that also has great potential for diagnostic purposes in clinical settings (Valdes et al., 2021; Zhao et al., 2017).

In recent years, various specialized ExM variants have been described, focusing on improving preservation of specific structures (Gambarotto et al., 2019), expanding tissues and entire organisms (Chen et al., 2021; Ku et al., 2016; Sim et al., 2022; Yu et al., 2020) including human biopsy tissues (Valdes et al., 2021; Zhao et al., 2017), and compatibility with other super-resolution modalities (Gao et al., 2020; Halpern et al., 2017; Shaib et al., 2022; Xu et al., 2019). Since the effective resolution of ExM is in part limited by the expansion factor, there has also been a push to develop higher expanding methods compared to the original ExM method either by varying the crosslinking concentration (Chen et al., 2015; Damstra et al., 2022), using alternative crosslinking chemistry (Li et al., 2022; Truckenbrodt et al., 2019), or using iterative approaches (Chang et al., 2017; M’Saad and Bewersdorf, 2020). The push for higher expansion and more complex ExM applications means that robust characterization of the exact expansion factor and of possible local deformations will be essential for biological reproducibility, quantitative accuracy, and diagnostic applications.

In most cases, the expansion factor of an imaged sample is determined macroscopically by measuring the size of the expanded gel. In addition, ExM variants are often validated for microscopic expansion factor using cellular reference structures, such as nuclear pore complexes (NPCs) or clathrin coated pits (CCPs) that have a distinct size known from other modalities such as electron microscopy. Alternatively, the precise local expansion factor can be determined by correlating a larger reference structure, e.g., a stained cell, before expansion to the same structure after expansion. This is a powerful approach as it can also be used to map local deformations that occur during expansion and sample mounting. However, due to the physical differences between pre- and post-expanded samples, robust correlation is time consuming, laborious, relies on efficient sample navigation and efficient labeling of the structure of interest, and is low throughput – making it challenging to repeatedly assess reproducibility and variability in terms of expansion and deformation for different gels and samples. Current attempts to directly measure the microscopic expansion factor without using reference structures or pre-expansion imaging are not readily compatible with biological samples and do not provide information about local deformations (Scheible and Tinnefeld, 2018). Therefore, easy to use reference-free quality control mechanisms are currently lacking, significantly hampering widescale adoption of ExM for quantitative purposes, including diagnostic applications.

Here, we introduce GelMap, a workflow wherein a gel-embedded reference grid is used to intrinsically calibrate hydrogels. GelMap functions as an expandable ruler that accurately reports (local) expansion factors, enables reference-free deformation mapping, and facilitates sample navigation. This reference grid can be directly used as a culturing substrate or be incorporated into the gel during later steps of sample preparation. By primarily using protein photolithography to pattern GelMap grids onto coverslips, we demonstrate that depending on the experiment, different grid designs, patterned proteins or fluorescent groups can be incorporated into the expansion hydrogel together with any biological specimen to reliably report expansion factors and enable computational correction of local deformations. Additionally, GelMap also solves the challenge of sample navigation in expansion microscopy by directly incorporating a fluorescent coordinate system into the ExM hydrogel. This facilitates experiments in which live-cell imaging is followed up by expansion microscopy. Finally, we demonstrate that GelMap grids can be used beyond cultured cells by imprinting them during gelation, enabling calibration and deformation correction of expanded brain tissue slices. In summary, GelMap provides a key quality control step for expansion microscopy, ready for easy adoption in existing established ExM workflows in both research and clinical settings.

## RESULTS

### Development of GelMap

Quantitative interpretation of expanded samples relies on robust characterization of expansion fidelity, which could be compromised by local anisotropies in expansion due to differences in gel composition, density, or mounting. We reasoned that ExM hydrogels could be intrinsically calibrated by incorporating a scalable fluorescent fiducial pattern, which would allow for precise determination of the local microscopic expansion factor. By generating a culturing surface that contains a fluorescent protein-based pattern of repeating squares (Fig. 1A, 1), and directly culturing cells on patterned coverslips, both the cells and the pattern would be incorporated into the ExM hydrogel simultaneously (Fig. 1A, 2). During expansion the pattern will expand together with the cells and act as a scalable ruler to determine the expansion factor. Since the marker is comprised of a repeating grid of known dimensions, local deformations that occur during expansion should be reflected in a visibly distorted grid (Fig. 1A, 3) that could be used to computationally correct for deformation without relying on a pre-expanded reference state (Fig. 1A, 4). To test this idea, we set out to develop a scalable fluorescent grid compatible with ExM chemistry that would be transferable from culture surface to hydrogel and biologically inert to ensure biological reproducibility and compatibility with all cell types.

**Figure 1:**
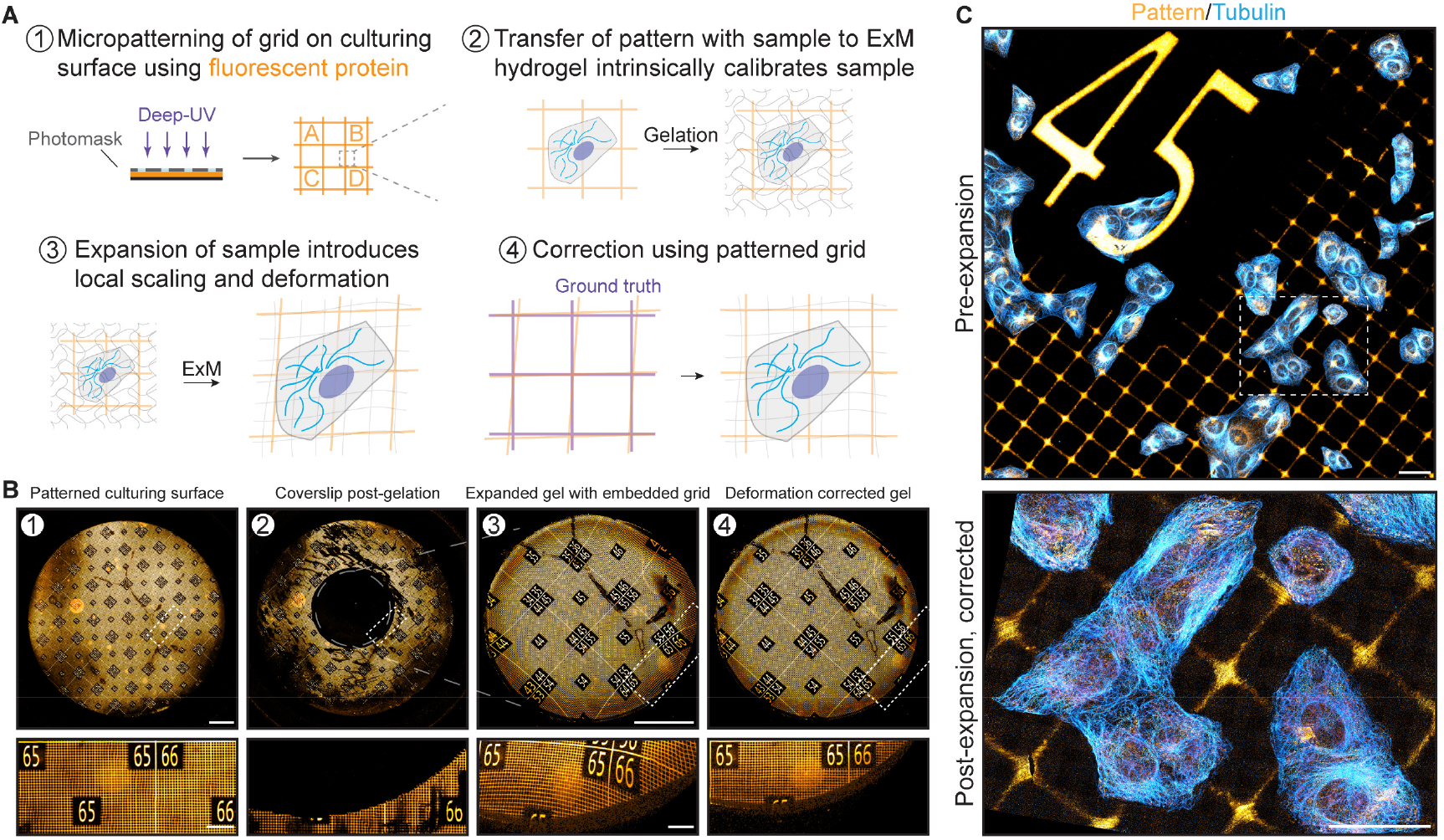
GelMap enables intrinsic calibration and deformation mapping in expanded hydrogels. **A)** Schematic of GelMap. **B)** Representative images of the indicated steps of GelMap. (1) 18 mm glass coverslip patterned with fluorescent laminin and amplified with antibody labeling (orange) shown before gelation. (2) The same coverslip after digestion of a locally polymerized gel using a silicone mold in the middle of the coverslip (6 mm diameter), showing complete transfer of protein pattern from the coverslip. (3) The polymerized and digested gel corresponding to the region indicated with grey circle in (2) was expanded 3x and imaged, showing anisotropic deformation. (4) The resulting acquisition was used for landmark based registration to the pre-expanded image in (1) for correction of local deformations in the gel. **C)** Top: U2OS cells stained for tubulin (cyan) cultured directly on laminin-patterned coverslip and imaged before expansion. Bottom: Region marked in top image after expansion (expansion factor: 9.6x) using TREx, corrected using GelMap. Scale bars: B: 2 mm, zoomed regions 400 µm. C: 40 µm. Scale bars in expanded samples reflect pre-expansion sizes.

We used protein photolithography with deep-UV to generate a fluorescent pattern as it is commonly used for biological applications, including micropatterning (Théry, 2010), has sufficient spatial resolution to pattern grids with micrometer scale, and is high throughput and easily scalable. We designed various chrome-quartz photomasks to test different numbered grid designs, with repeated features ranging in size from 10 µm to 40 µm to focus on mapping deformations, and larger grid sizes from 200 µm to 400 µm that included markers to aid sample navigation (Fig. S1A, B). We tested our patterning approach using coverslips coated with laminin, an extracellular matrix protein (ECM) conjugated to a fluorophore and found we could successfully pattern entire coverslips (Fig. 1B, 1), with multiple designs (Fig. S1C). Next, we tested whether the fluorescent pattern could be transferred from the culturing substrate to a hydrogel by gelating and expanding a region in the middle of the coverslip using Ten-fold Robust Expansion (TREx) microscopy (Damstra et al., 2022). Imaging of the same coverslip after gelation confirmed the pattern was efficiently transferred from the substrate to the hydrogel (Fig 1B, 2). When the resulting gel was imaged after moderate expansion (3x) to facilitate imaging of the entire gel, we observed local deformations that distorted the uniformity of the pattern and were particularly pronounced near the edge of the gel (Fig. 1B, 3). By registering the deformed image to the pre-expansion pattern, the deformations could be corrected using non-linear thin plate spline transformation of the deformed image as shown previously (Chen et al., 2015; Damstra et al., 2022; Jurriens et al., 2020), which restored the uniformity of the pattern in the expanded gel (Fig. 1B, 4). Finally, we tested whether our approach could be directly used as a culturing substrate by sterilizing a grid and growing cells directly on the protein patterned coverslip. Following fixation and immunostaining for tubulin and laminin to amplify the fluorescent grid, the sample was imaged pre-expansion and subsequently expanded. Pre-expansion imaging confirmed that cells grew irrespective of the underlying protein grid (Fig. 1C, pre-expansion). Comparison of pre- and post-expansion images confirmed that cells and the pattern are expanded together, and by aligning the expanded image to a virtual reference grid local deformations could be corrected for (Fig. 1C).

In the described workflow, the pattern was amplified using antibody labeling (Fig. S1D). Using the laminin pattern, we observed a small degree of labeling of cell-derived laminin when amplifying the pattern, which could hinder grid identification and deformation mapping. Therefore, we tested patterning efficiency using fibrinogen, a different ECM-protein, as well as a nanobody against a non-biological target (R2-myc-his, hereafter NBD) that contains an orthogonal myc-tag that can be used for amplification without intracellular background (Fig. S1E). Following conjugation to a fluorophore, we found the bulkier ECM proteins patterned as efficiently as the smaller globular nanobody, and in both cases the sterilized patterns could be used as a culturing surface.

Because some applications of ExM (e.g. expansion of tissue samples) do not involve a culturing surface or could require calibration and correction in all three-dimensions, we reasoned that an approach to create such grid maps within a polymerized, non-expanded gel could carry advantages. To that end, we introduced an acrylate-modified photoactivatable rhodamine into the ExM gelation solution. After polymerization of the hydrogel, we introduced a pattern into the pre-expanded hydrogel by local photo-uncaging using targeted illumination. These photoactivated patterns were stable and expanded along with the ExM hydrogel (Fig. S1F), providing an alternative implementation of GelMap that could in principle be extended to three-dimensions.

Thus, our approach provides an easy-to-use, flexible method to intrinsically calibrate ExM hydrogels that is compatible with biological specimen and ExM chemistry. Because this approach provides quantitative scaling information, reports on local deformations and can be used for easy navigation, we termed the resulting method GelMap.

### Correction of expansion anisotropy using GelMap

We next used GelMap to assess the expansion inhomogeneities and deformations that occur during expansion. We first examined how the estimated macroscopic expansion factor, as measured by the size of the expanded gel, relates to the measured microscopic expansion factor. We expanded GelMap grids without cells using TREx and compared the microscopic expansion factor of individual squares (2 regions, 849 individual squares, expansion factor 9.0 ± 0.2 (mean ± SD)) with 7 individual estimates of macroscopic expansion factor by unbiased participants. While the average measurement from the participants was consistent with the calculated average of the true expansion factor (9.0 ± 0.3, mean ± SD), the individual estimates ranged from 8.6 to 9.4. This highlights possible errors that can be introduced from individual measurements (Fig. S2A). We next looked at the deformation of individual 40 × 40 µm squares within the same gel. We defined the squareness of each square within the field of view as a metric for ExM fidelity (accounting for both stretching and skewness) and observed significant variability across multiple gels and regions (3 gels, 3 regions per gel, 3574 individual squares), (Fig. S2B). We could observe some regions that were hardly deformed (Fig. 2A, region 1), and other regions where very strong deformations were apparent, for example near tears (Fig. S2C). We also noted more subtle anisotropies that became visible using GelMap, which may remain unrecognized when looking at biological structures alone (Fig. 2A, region 2). Confirming these qualitative observations, plotting the distribution of squareness values for both regions revealed markedly more deformation in region 2 (Fig. 2B). Together, these experiments underscore the need for local quality control mechanisms to standardize quantitative measurements in ExM.

**Figure 2:**
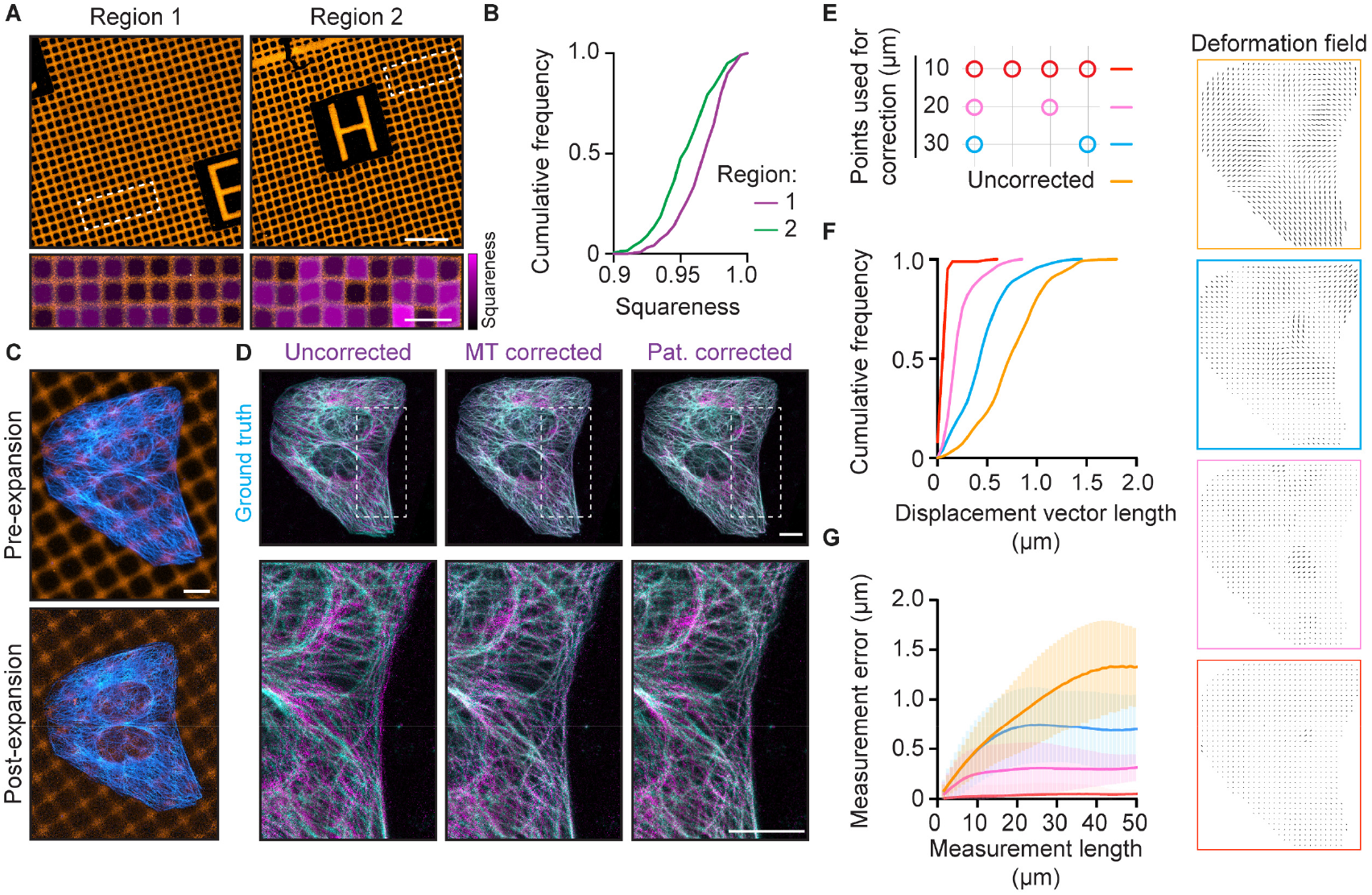
Validation of GelMap-based corrections using pre- and post-expansion imaging. **A)** Two representative regions of expanded GelMap grids (20 µm squares) from the same gel with deformation as defined by squareness color-coded in magenta in zooms of boxed regions (expansion factor: 9.0 and 8.7 for region 1 and 2, respectively). **B)** Cumulative frequency distribution of squareness values of individual squares in region 1 (n=453, magenta) and region 2 (n=396, magenta) from (A). **C)** Pre- and post-expansion images of U2OS cells cultured on nanobody-patterned GelMap grids (grid size: 10 µm, expansion factor: 9.1), stained for tubulin (cyan) and myc-tag (orange). **D)** Linear (left column; uncorrected) and non-linear transformation (middle column; MT corrected) of the post-expansion tubulin channel registered to the pre-expanded ground truth tubulin channel, overlayed with pre-expanded ground truth image. Pattern-corrected tubulin channel (10 µm grid spacing) overlayed with the pre-expanded ground truth image (right column, pat. corrected). Boxed regions correspond to zooms below. **E)** Schematic of landmark spacing used for pattern correction. The 10 µm GelMap grid was used for landmark registration, with landmarks removed as indicated to mimic GelMap grids with larger spacing. **F)** Quantification of displacement vector length for equidistantly spaced points (1 µm) in the cellular area for pattern-corrected images registered to ground truth tubulin channel. Corresponding deformation fields are color-coded and shown on the right. **G)** Quantification of measurement error (deviation from expected distance assuming isotropic expansion) between pairs of points for a measurement length (mean ± SD). Scale bars: A: 100 µm, zoomed regions 40 µm. C-D: 10 µm

We next explored the effectiveness of GelMap for correcting deformations on the cellular scale. We cultured cells on an NBD-patterned coverslip, fixed and stained for tubulin, amplified the pattern using the orthogonal myc-tag, and then examined the corrective power for different grid sizes compared to correction using the pre-expansion image of the tubulin (MT) channel (Fig. 2C-D). First, we visualized the overall deformation by registering and overlaying the pre- and post-expansion tubulin channels using a landmark-based linear similarity transformation (Fig. 2D, uncorrected), which was subsequently corrected using non-linear thin plate spline transformation (Fig. 2D, MT corrected). Next, we used GelMap and first registered the pre- and post-expansion grid images using non-linear transformation, which was then used to correct the expanded tubulin channel (Fig. 2D, pattern corrected).

To quantify the corrective power of GelMap, we first generated three pattern-corrected images, for which we varied the number of landmarks used for registration to represent grid dimensions of 10 × 10 µm, 20 × 20 µm and 30 × 30 µm (Fig. 2E). To determine the residual errors in these GelMap-corrected images, we then registered these images to the pre-expansion tubulin image. By comparing linear and non-linear transformations, we could then calculate for each used grid dimension a displacement vector map that shows the remaining deformation for equidistantly spaced points (1 µm) within the cellular area (Jurriens et al., 2020). Inspection of the distribution of vector lengths revealed that finer grids resulted in less residual errors (Fig. 2F, S2D-F). The average residual error decreased from 0.76 ± 0.3 µm (mean ± SD) without correction to 0.07 ± 0.05 µm (mean ± SD) for correction every 10 µm.

An alternative measure for deformation is to compare measurement lengths between pairs of points after expansion to the expected distance in the case of uniform expansion, and plot the average fractional deviation as a function of the measurement length (Chen et al., 2015) (Fig. 2G, S2G). Again, we observed that correction using grids with smaller spacing resulted in smaller errors for a given measurement length. While the uncorrected curve saturates at a measurement length of ∼45 µm, the curves for measurements after correction saturate around 25 µm, 15 µm and 7 µm for correction using landmarks spaced 30 µm, 20 µm, and 10 µm, respectively.

These experiments have two important implications. First, while all grid sizes can correct for deformation, the absolute corrective power of GelMap is inversely correlated with the grid feature size. Second, as GelMap is effective in correcting deformations via registration to the pre-expanded dimensions, registration to a virtual reference grid of identical dimensions must be equally effective. This in turn negates the need for pre-expansion image acquisition. Therefore, by intrinsically calibrating the ExM hydrogel with a fluorescent grid that scales with the expansion factor and deforms with anisotropy, we have developed a robust quality control method for ExM.

### GelMap facilitates sample navigation in correlative live-ExM experiments

Following up live-cell microscopy experiments with expansion microscopy is a promising approach that can provide novel insights into the nanoscale structures that underly cellular dynamics. However, the sample processing steps required after live-cell imaging makes it challenging to locate back specific imaged cells in the expanded gels. This is particularly the case for high-expanding gel recipes, as the volume that contains the cell of interest increases with the expansion factor cubed. To address this challenge, we added a coordinate system into our GelMap grids. This would enable us to image a cell, locate the position of the cell on the grid (Fig. 3A, 1), fix and stain the cells for the structure of interest, gelate the sample, and excise the gel region that contains the cell (Fig. 3A, 2). The excised gel fragment could then be expanded, after which the grid facilitates correlation of the expanded cell with the live-cell acquisition, as well as analysis of local deformations and expansion factor (Fig. 3A, 3).

**Figure 3:**
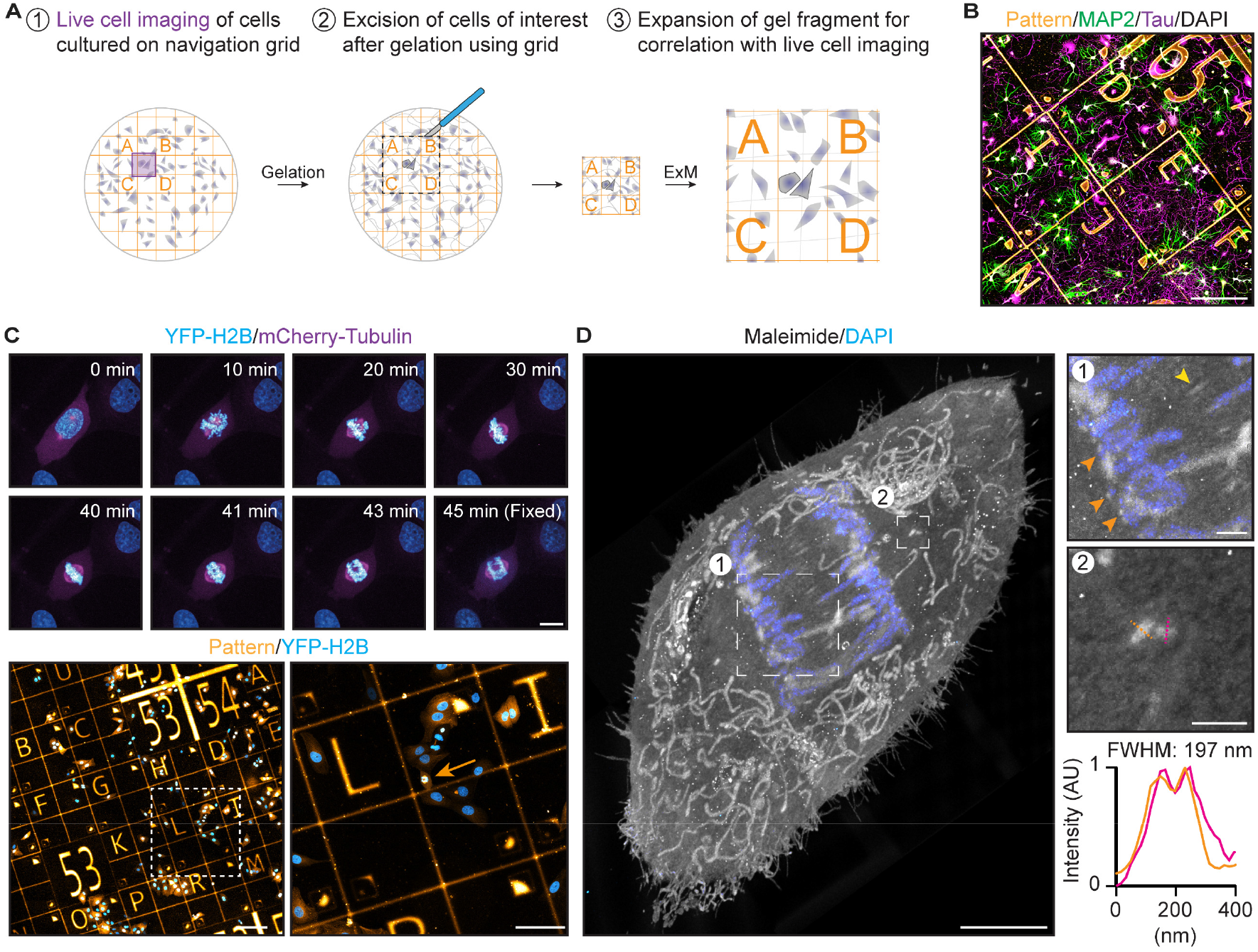
GelMap facilitates sample navigation for correlative live and expansion microscopy. **A)** Schematic of workflow for correlative live and expansion microscopy (1) Cells of interest cultured on GelMap coverslips are followed live to image a dynamic process. (2) Cells are incorporated into the ExM hydrogel, digested, and the region of interest as indicated by the pattern is excised before expansion. (3) Gel fragment is expanded, and pattern is used to find back the cell of interest for correlation. **B)** Dissociated neurons cultured on GelMap navigation grid, stained for MAP2 (green), Tau (magenta), DAPI (gray), and myc-tag (orange). **C)** Time-lapse imaging of a mitotic cell stably expressing YFP-H2B and mCherry-Tubulin. During anaphase, the cell was fixed on the microscope stage. The cell was located on navigation grids and expanded using TREx. **D)** Maximum projection of expanded cell from time-lapse imaging in (C) stained for total protein using maleimide and DAPI (expansion factor: 10.7). Zooms are indicated by boxed regions. General protein stain reveals several ultrastructural features including cell morphology, intracellular organelles such as mitochondria, the spindle midzone (yellow arrow), presumptive kinetochores (zoom 1, orange arrows) and centrioles (zoom 2). Line scan and quantification of centriole width indicated in orange and magenta. Scale bars: B: 200 µm, C: timelapse 10 µm, below: 200 µm (left) and 100 µm (right), D: 5 µm, zoomed regions 1 µm.

We employed various navigation grids (Fig. S1B) and tested compatibility of GelMap with a top-coating of substrate to facilitate live imaging of sensitive cells such as neurons, confirming neuronal growth and morphology are unperturbed (Fig. 3B). To demonstrate correlative expansion microscopy, we cultured cells stably expressing YFP-H2B and mCherry-tubulin on the navigation grid and synchronized the population using thymidine. We picked a cell of interest and followed its progression through mitosis. During anaphase, the cell was rapidly fixed on the microscope stage and located on the navigation grid (Fig. 3C, Video S1). We next expanded the coverslip using TREx, excised the region of interest (relying on navigation provided by GelMap and stained for total protein using maleimide. In the expanded sample, we could use the ultrastructural context provided by maleimide to visualize the cell morphology, organelle distribution, and key components of the mitotic spindle, including the spindle midzone, the chromosomes and presumptive kinetochores, and both centrioles on either side of the spindle (Fig. 3D, S3B, Video S1). Furthermore, by registering the grid to the ground truth we could extract the exact expansion factor for the gel (Fig. S3A) and determined the diameter of the centrioles in the cell to be 197 nm, consistent with previously reported values using ExM (Gambarotto et al., 2019).

### GelMap imprinting onto tissue slices during sample processing

One of the key advantages of ExM is that it can be used to directly expand tissues and organisms (Chen et al., 2015, 2021; Ku et al., 2016; Sim et al., 2022; Yu et al., 2020). We therefore tested whether GelMap could be applied to non-adherent biological samples by incorporating the grid at the stage of sample preparation. Gelation is often performed in a gelation chamber that is closed off during polymerization. We therefore set out to expand mouse brain tissue slices and used a patterned coverslip to close off the gelation chamber before gelation, ensuring contact between the tissue slice and the patterned coverslip (Fig. 4A). Indeed, this procedure results in reliable imprinting of the GelMap grid on one side of the tissue (Fig. 4B). Following expansion and staining of the tissue with a NHS general protein stain, we visualized the imprinted fluorescent pattern in combination with the tissue. This revealed significant distortions of the grid that indicated local deformations during expansion. These distortions could be subsequently corrected by aligning the expanded image to a virtual reference grid followed by non-linear transformation (Fig. 4B).

**Figure 4:**
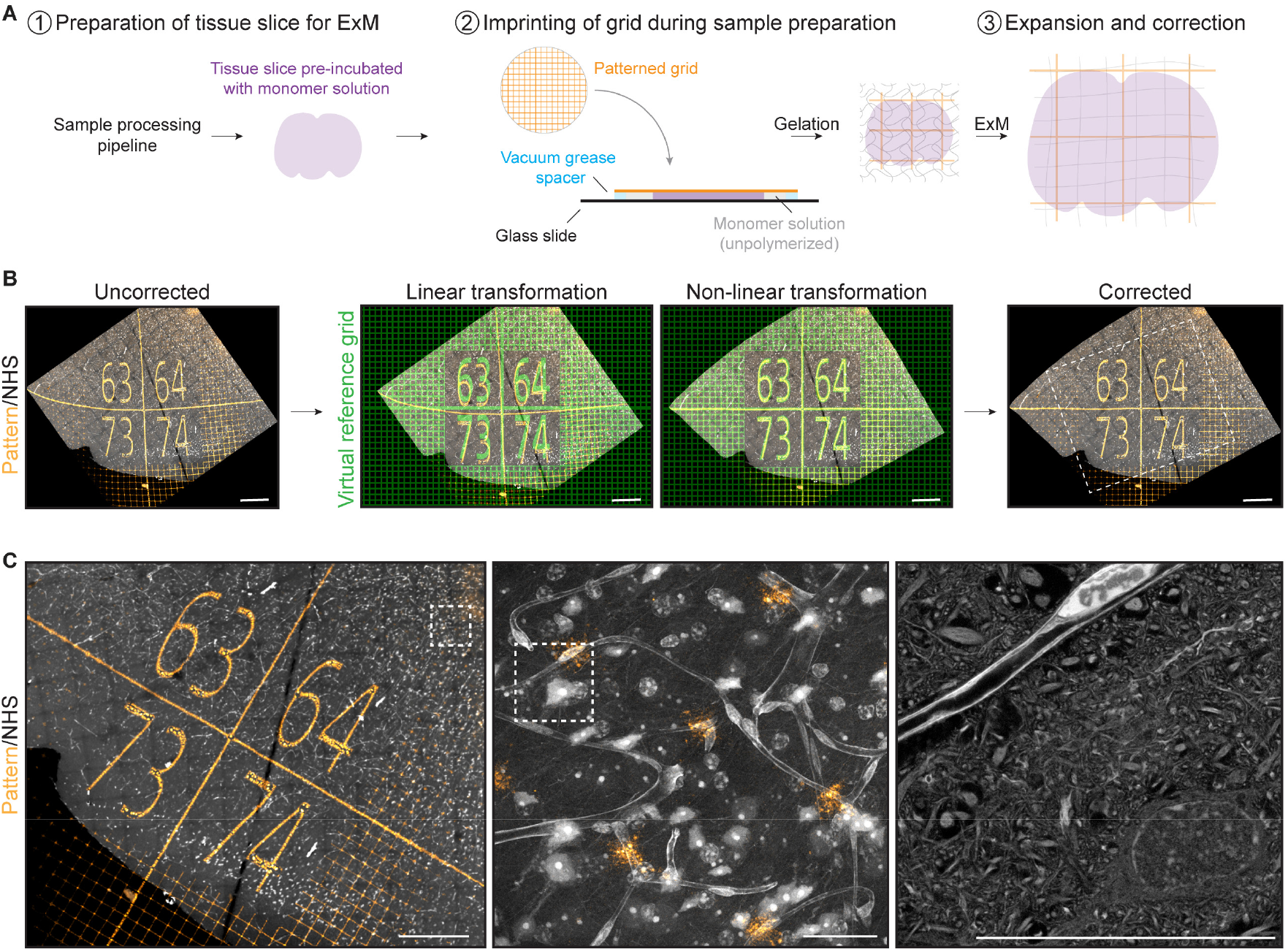
GelMap can be imprinted onto tissue slices and used for correction. **A)** Schematic of workflow when using GelMap for tissue slices during sample preparation. **B)** Region of dissected mouse brain tissue (cortex) expanded using TREx (expansion factor: 7.6) and stained for total protein (gray) and myc-tag (orange) and imaged by confocal microscopy (left). The pattern channel was registered to a virtual reference grid (green) to correct for deformation (middle and right panel). The resulting non-linear transformed protein channel was used for subsequent visualization of corrected tissue organization. **C)** Using GelMap to bridge scales of tissue vasculature organization from 1.3×1.2 mm (left) to 110×110 µm and 25×25 µm (corrected dimensions) high resolution zooms of blood vessel with surrounding tissue. Zoomed regions indicated with white boxes. Scale bars: B: 200 µm, C: 200 µm (left), 20 µm (middle and right).

Using GelMap, we could bridge multiple layers of tissue organization; from a low-resolution overview tilescan (1.3 × 1.2 mm, corrected dimensions), to a higher resolution overview of the local arrangement of cells and tissue vasculature (110 × 110 µm, corrected dimensions), down to high resolution ultrastructural details of neuronal somata, surrounding neuropil, and the vascular endothelium (25 × 25 µm, corrected dimensions) (Fig. 4C). From the GelMap grid the exact expansion factor of the tissue can be extracted, which is essential for precise quantitative measurements. These results demonstrate that the GelMap approach can be added to various existing ExM sample processing pipelines to provide intrinsic calibration and deformation mapping for expanded tissue.

## DISCUSSION

Here, we introduce GelMap as an approach to intrinsically calibrate ExM hydrogels through the incorporation of a fluorescent grid that scales with the expansion factor and deforms upon anisotropic expansion. Our approach overcomes three key challenges of ExM. First, it provides a robust way to determine the exact expansion factor in relevant regions of interest. Second, it enables one to routinely quantify and correct local deformations without the need for a pre-expanded reference image. Third, it encodes a fluorescent coordinate system to the hydrogel that aids sample navigation of expanded samples. Importantly, GelMap can be easily incorporated into existing ExM workflows to provide a standardized quality control step.

While the use of pre-patterned substrate-attached proteins to create two-dimensional grids is the most straightforward implementation of GelMap, we have also shown that grids can be created by structured photoconversion of gel-incorporated fluorophores before expansion. This opens the way for creating three-dimensional grids, for example in combination with two-photon conversion to ensure localized conversion in the third dimension.

GelMap provides a simple and robust method to calibrate expanded gels, to map and correct expansion-induced deformations, and to facilitate sample navigation in correlative microscopy approaches. We anticipate that this will aid the development and validation of new gelation and expansion protocols and facilitate the widespread implementation of Expansion Microscopy into imaging applications that require robust and reliable calibration, both in basic science and in clinical applications.

## ACKNOWLEDGEMENTS

We thank Wilco Nijenhuis for helpful advice and Eugene Katrukha for helpful discussions about image analysis. This work was supported by EMBO long-term fellowship (EMBO ALTF 407-2017), the European Research Council (ERC Consolidator Grant 819219), and the Eindhoven, Wageningen Utrecht Alliance through the Centre for Living Technologies.

## COMPETING INTERESTS

H.G.J.D., J.B.P. and L.C.K have filed a patent application covering the presented methods.

## MATERIALS AND METHODS

### Photomask design

CleWin 5 (WieWeb) software was used for design of chrome-quartz photomasks. The following base design was used: 2 × 2 cm area consisting of 100 (00-99) 2 × 2 mm tiles, containing five 400 × 400 µm numbers (Fig. S1A). Variants on the base design followed either a repeated grid pattern for use in mapping gel deformation, or a coordinate letter system to further aid in sample navigation (Fig. S1B). Photomasks were fabricated by Toppan (Toppan photomasks Germany GmbH), with features (numbers, lines) chrome and blank spaces quartz.

### Conjugation

Conjugation of protein or nanobody was performed with an 8-fold (Laminin; Roche) or 4-fold (R2-myc-his; a gift from S. Oliveira) molar excess of NHS-ester (ATTO 425, ATTO 647N; ATTO-TEC GmbH, Alexa 488, Alexa 594; Sigma Aldrich) in a minimal volume of MQ. Protein/nanobody and NHS-ester were mixed by vortexing and incubated overnight at 4 °C before storing 10 µg aliquots at -20 °C. Fibrinogen pre-conjugated with Alexa Fluor 647 (Invitrogen) was stored in sodium bicarbonate buffer, pH 8.3.

### Preparation of patterned coverslips

Coverslips (Ø18 mm, #1.5; Marienfeld, 107032) were washed for 10 min in acetone, sonicated for 20 min in 50 % methanol and sonicated for a further 20 min in 0.5 M KOH before washing three times in MQ. Coverslips were then washed in 100% ethanol and dried under a flow of nitrogen before storage. Cleaned coverslips were activated with air plasma (PDC-002, Harrick Plasma) for 1 min before incubation with 2.5 µg fluorescent protein in 100 µl MQ per coverslip for 1 h at room temperature. Coverslips were dried at room temperature on a clean tissue for 30 min before being exposed to deep-UV (∼250 nm) through a micropatterned chrome/quartz photomask for 4 min, using an UVO cleaner 42-220 (Jelight Company Inc.). Patterned coverslips were washed with PBS for storage at 4 °C. Photomasks were cleaned by 10 min acetone and isopropanol alcohol washes, before rinsing in MQ and drying under a flow of nitrogen. Photomasks were exposed to air plasma for 20 min prior to patterning.

Prior to cell seeding, patterned coverslips were sterilised with 70 % ethanol and washed three times in sterile PBS. For neuronal experiments, patterned coverslips were top-coated with poly-L-lysine (37.5 μg/ml) and laminin (1.25 μg/ml) in 0.1 M Borate Buffer, pH 8.5. Laminin patterned coverslips were therefore not used for neuronal experiments.

### Photo-uncageable rhodamine patterning within polymerized hydrogels

To generate a photoactivatable molecule that could be covalently incorporated into polymerized hydrogels, a two-step conjugation was performed of NVOC2-Q-rhodamine-5-PEG3-azide (Sigma-Aldrich, 768693) to acryloyl-X SE (AcX) (Thermo Fisher, A20770) using a sulfo-DBCO-amine linker (Broadpharm BP-23309). First, Sulfo-DBCO-amine (40 mM) was reacted with AcX (21 mM) in DMSO for 1 h at RT to form DBCO-AcX. Next, a 2-4x molar excess of NVOC2-Q-rhodamine-5-PEG3-azide was reacted with DBCO-AcX for 1 h at RT. By mixing the thus formed molecule into the unpolymerized gel solution at a final concentration of 120 μM, the dye becomes covalently linked to the ExM hydrogel during gelation. Using local UV illumination, photoactivated patterns could be generated, expanded with the hydrogel, and used for intrinsic calibration.

### Cell culture and cell synchronization

U2OS (ATCC) cells were cultured in DMEM medium supplemented with 9 % Fetal Bovine Serum and 1 % penicillin/streptomycin (GIBCO). Primary hippocampal neurons were maintained in Neurobasal medium supplemented with 1 % B27 (GIBCO), 0.5 mM glutamine (GIBCO), 15.6 μM glutamate (Sigma), and 1 % penicillin/streptomycin (GIBCO). For imaging of mitosis, U2OS cells expressing YFP-H2B and mCherry-tubulin (a gift from W. Nijenhuis) were synchronized using a double thymidine block (Chen and Deng, 2018). In short, cells were seeded at 25 % confluency, 16 h after seeding incubated with 2 mM thymidine for 24 h, incubated with complete medium for 9 h, blocked again using 2 mM thymidine for 29 h, and finally released from second block in complete medium 12 h prior to imaging.

### Fixation and immunostaining

For correlative deformation mapping experiments, cells were extracted using pre-warmed 0.35 % Triton X-100 + 0.2 % glutaraldehyde in MRB80 for 1 min, followed by 4 % paraformaldehyde fixation in PBS for 10 min. For experiments using protein stains, cells were fixed with prewarmed (37 °C) 4 % paraformaldehyde + 0.1 % glutaraldehyde + 4 % sucrose in PBS for 10 min. Next, cells were washed with PBS and permeabilized with PBS + 0.2 % Triton X-100. Blocking and antibody labeling steps were performed with 3 % bovine serum albumin in PBS. The patterned grids were amplified by immunostaining for laminin, fibrinogen, or myc-tag (R2-myc-his) along with antibody labelling for specific structures. The following primary antibodies were used in this work: rabbit anti-laminin (1:100, Abcam ab11575), rabbit anti-myc-tag (1:100, Cell Signalling Technology 2272), rabbit anti-fibrinogen (1:100, Abcam ab34269), rat anti-tubulin YL1/2 (1:200, Abcam ab6160), mouse anti-tau (1:500, Sigma-Aldrich MAB3420), chicken anti-MAP2 (1:500, Abcam ab5392). The following secondary antibodies were used in this work: goat anti-rabbit IgG Alexa Fluor 594 (Invitrogen A-11037), goat anti-rabbit IgG Alexa Fluor 488 (Invitrogen A-11029), goat anti-rat IgG Alexa Fluor 488 (Invitrogen A-11006). goat anti-chicken IgY DyLight 488 (Invitrogen SA5-10070), goat anti-mouse IgG Alexa Fluor 647 (Invitrogen A-21236), all at a 1:250 dilution.

### Tissue fixation and sectioning

All animal experiments were carried out according to the regulations of Utrecht University and in agreement with Dutch law (Wet op de Dierproeven, 1996) and European regulations (Directive 2010/63/EU). 10-month-old, TRAP2 heterozygous mice (Jax #030323) were transcardially perfused with ice-cold fixative solution (4 % formaldehyde and 20 % acrylamide in PBS, pH 7.4). Brains were removed and post-fixed in fixative solution overnight at 4 °C. Fixed brains were washed three times for one hour in PBS at RT and cut coronally into 100 µm thick sections using a vibratome (Leica VT1000 S). After cutting, fixed brain sections were stored in PBS at 4 °C.

### Tenfold Robust Expansion (TREx) Microscopy

Tenfold Robust Expansion (TREx) microscopy was performed according as previously published (Damstra et al., 2022). Cells were treated with 100 µg/ml acryloyl-X SE (AcX) (Thermo Fisher, A20770) in PBS overnight at RT. TREx gelation solution was prepared containing 1.1 M sodium acrylate, 2.0 M acrylamide (AA), 50 ppm N,N’-methylenebisacrylamide (bis), PBS (1x), 0.15 % APS, 0.15 % TEMED. Gels were prepared in a gelation chamber consisting of a parafilm covered glass slide with a silicone gasket (Sigma-Aldrich, GBL66410). Gels were left to gelate for 1 h at 37 °C. Next, samples were transferred to a 12-well plate and digested with 7.5 U/ml Proteinase-K (Thermo Fisher, EO0491) in TAE buffer (containing 40 mM Tris, 20 mM acetic acid and 1 mM EDTA) supplemented with 0.5 % Triton X-100, 0.8 M guanidine-HCl, and DAPI for 4 hours at 37 °C. For correlative experiments, regions of interest were located after digestion using fluorescent grid, excised, and expanded using an EVOS imaging system equipped with Plan Fluor 10x/0.3 objective (ThermoFisher Scientific). The gel was transferred to a Petri dish, water was exchanged 2x 30 minutes and the sample was left in MQ to expand overnight. Prior to imaging the cells were trimmed using a scalpel blade to fit in an Attofluor Cell Chamber (Molecular probes A-7816), or a custom designed 3D printed imaging chamber. For samples stained for total protein using maleimide, gels were washed 2x 15 min in PBS after gelation and incubated with 20 μg/mL Atto 647N maleimide (Atto-Tec, AD 647N) in PBS prepared from a 20 mg/mL stock solution in DMSO for 1.5 h at RT with shaking. Next, samples were rinsed in PBS, digested, and expanded as described above.

### Expansion of tissue and incorporating patterned grids during sample processing

For expansion of tissue, brain slices were pre-incubated with TREx gelation solution with 15 µg/mL 4-hydroxy-TEMPO added to delay premature gelation for 30 min on ice. To create the gelation chamber, tissue slices were laid out on a microscopy slide with 4 dabs of vacuum grease surrounding the slice. A GelMap coverslip was placed with the protein grid facing the tissue on top of the dabs of vacuum grease and pressed down ensuring contact with the tissue slice. The gelation chamber was filled with gelation solution from the side and transferred to 37 °C for 1 h. After gelation, the gel surrounding the tissue was trimmed and the sample was disrupted for 3 h at 80 °C in disruption buffer containing 5 % SDS, 200 mM NaCl, and 50 mM Tris pH 7.4. After disruption, gels were washed in PBS and stained with 30 µg/mL NHS-ester conjugated to ATTO488 (ATTO-TEC GmbH) for 2 h at RT. After staining, gels were washed and expanded in MQ. Prior to imaging the cells were trimmed using a scalpel blade to fit in a Attofluor Cell Chamber (Molecular probes A-7816), or a custom designed 3D printed imaging chamber.

### Imaging acquisition

ExM and pre-expansion images were acquired using a Leica TCS SP8 STED 3X microscope equipped with HC PL APO 20×/0.75 dry and HC PL APO 86×/1.20W motCORR STED (Leica 15506333) water objectives. A pulsed white laser (80 MHz) and a 405 nm DMOD Flexible UV laser were used for excitation. The internal Leica GaAsP HyD hybrid detectors were used with a time gate of 1 ≤ tg ≤ 6 ns. The set-up was controlled using LAS X. For photo-uncaging of rhodamine to create patterns in polymerized hydrogel, 50 × 50 µm field of views were scanned with 405 nm laser light to create a boxed pattern.

For imaging of pre-expanded neurons cultured on patterned coverslips, a Zeiss LSM 700 confocal setup consisting of an AxioObserver Z1 microscope with a Plan-Apochromat 20×/0.8 dry objective was used. The set-up was controlled using ZEN.

For live-cell imaging of mitosis, coverslips were mounted in complete medium and images acquired using a 60× (Plan Apo VC, NA 1.4; Nikon) oil-immersion objective on a Spinning Disc (Yokogawa CSU-X1-A1) Nikon Eclipse Ti microscope with Perfect Focus System equipped with a sample incubator (Tokai-Hit) and an Evolve 512 EMCCD camera (Photometrics), controlled with MetaMorph 7.7 software (Molecular Devices). Cobolt Calypso 491 nm and Cobolt Jive 561 nm lasers were used for excitation. Images were acquired every 60 s and fixed on the stage during imaging with pre-warmed fixative.

For imaging the chrome-quartz photomask and validating antibody amplification, an AMG EVOS digital inverted microscope was used, equipped with AMG 4×/0.13 Plan LWD PH and AMG 10×/0.3 Plan FL objectives.

### Image analysis

All imaging processing was done using FIJI (Schindelin et al., 2012). For non-linear transformation of post-expanded images using landmark-based registration of expanded samples to a (virtual) reference grid the plugin BigWarp (Bogovic et al., 2016) was used. All scale bars in figures represent pre-expansion dimensions, following correction for expansion factor using GelMap. Estimation of macroscopic expansion factor by unbiased participants was obtained by measuring the size of the expanded gel with a ruler and comparing it to the size of the original silicone mold.

For deformation correction using GelMap, regular landmarks provided by the grid were registered to either the pre-expanded grid or a virtual reference grid followed by non-linear thin plate spline transformation. For Fig. 2E-G and Fig. S2E-G, the total deformation was determined by landmark-based registration of pre- and post-expansion images of the microtubule cytoskeleton. The pattern correction was performed as described above and the residual error was quantified by registering the corrected image of the MT cytoskeleton to the pre-expansion ground truth. By comparing linear similarity transformation to non-linear thin plate spline transformation, the absolute deformation for equidistantly spaced points (1 µm) within the cellular area was calculated (Jurriens et al., 2020). Comparing measurement lengths between pairs of points after expansion to the expected distance in the case of uniform expansion and plotting of the average fractional deviation as a function of the measurement length was performed as previously described (Damstra et al., 2022). In all corrected post-expansion images, the expansion factor was determined from the BigWarp transformation (Jurriens et al., 2020).

Expansion anisotropy was quantified using a semi-automatic ‘squareness’ measurement of deformation. This measure was defined by the detection of the corners of each square in the GelMap grid, followed by determination of the length of the square diagonals as well as thevintersection angles of these diagonals. Squareness was then determined as the average of two metrics; (1) the ratio between the shorter and the longer diagonal length, and (2) the ratio between the smaller and the larger intersection angles of the diagonals. This metric takes into account both stretching and skewness and will yield a value between 0 and 1, with 1 corresponding to a perfect square and 0 corresponding to a completely flattened square, where two of the opposite corners are overlapping. The expansion factor was calculated for each square by dividing the measured area by the known pre-expansion area.

**Supplementary Figure 1:**
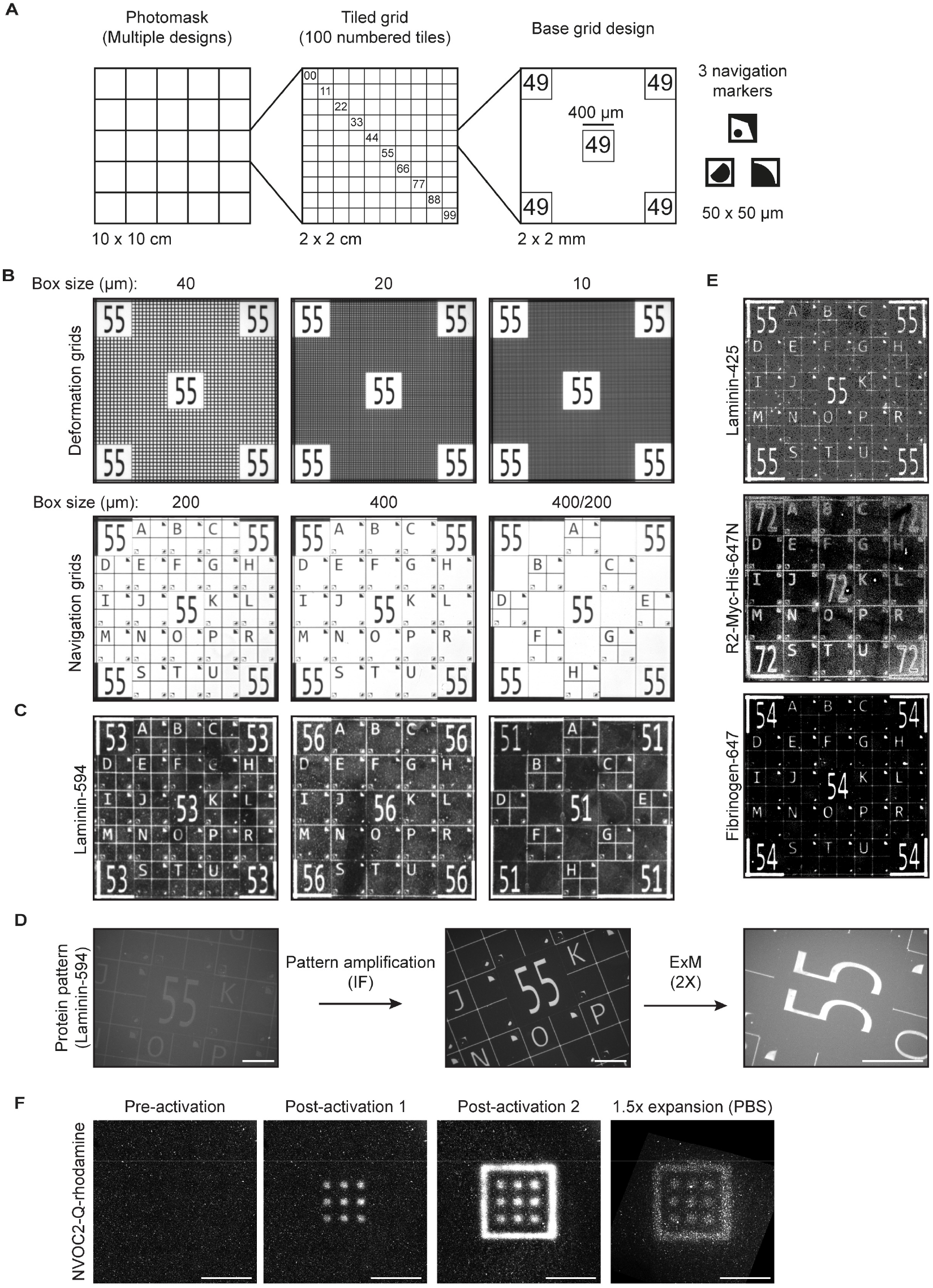
GelMap pattern design and validation. **A)** Photomask design showing tiled grid and the base grid design that can be adapted depending on the experiment. **B)** Widefield images of the photomask containing various grid designs focused on deformation mapping (top) and navigation (bottom). **C)** Laminin conjugated to Alexa-594 was patterned onto a glass coverslip using photolithography using various photomask designs. **D)** Widefield images of a laminin-594 patterned coverslip after patterning (left), after amplification of the signal using antibody labeling against laminin (middle) and moderately expanding using TREx (right). **E)** GelMap coverslips can be produced using multiple proteins (laminin, nanobody (R2-Myc-His) and fibrinogen) and various conjugated dyes. **F)** Alternative implementation of GelMap: an acrylate-modified photo-uncageable rhodamine was incorporated during gelation. Using targeted illumination, a fluorescent pattern was generated into the polymerized hydrogel. Finally, the pattern remains stable in the hydrogel and could be moderately expanded (expansion factor: 1.5) and aligned to pre-expanded dimensions for correction. Scale bars: base grid design: 2×2 mm, D: 200 µm, F: 20 µm.

**Supplementary Figure 2:**
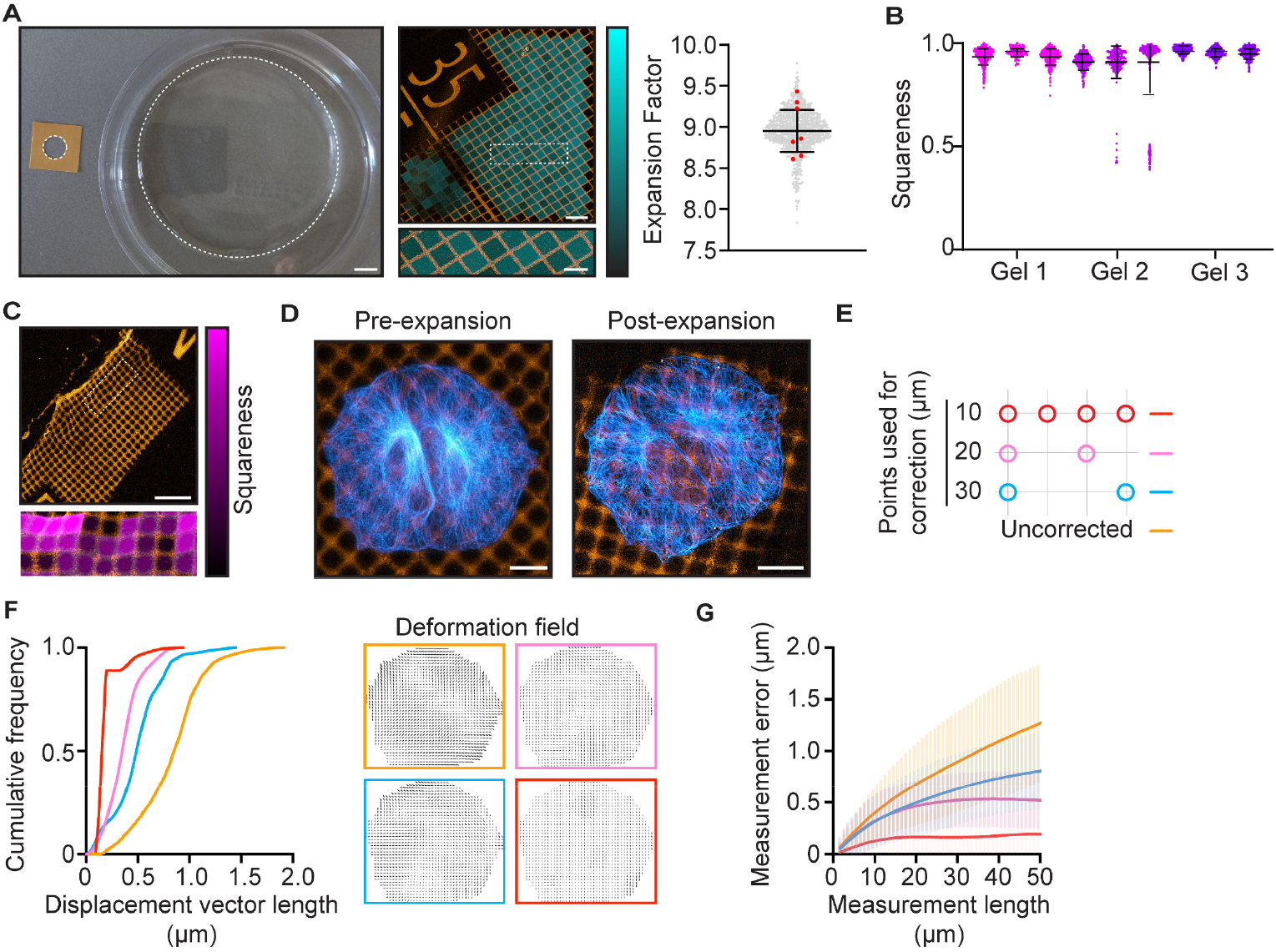
ExM anisotropy and GelMap validation. **A)** Left: example of typical macroscopic expansion factor determination with silicon mold on left and expanded gel on the right. Middle: expanded region showing GelMap grids (orange) from gel in (A), with the local expansion factor for each square color-coded and overlayed. Right: quantification of expansion factor of individual squares (2 regions, n = 849, expansion factor 8.9 ± 0.2 (mean ± SD), grey), with macroscopic measurements from unbiased participants overlayed (n = 7, expansion factor 8.9 ± 0.3 (mean ± SD), red). **B)** Quantification of squareness of individual squares for 3 gels, 3 regions per gel, n = 533, 116, 391, 383, 269, 756, 453, 396, 277, respectively, with mean ± SD squareness values of 0.94 ± 0.04, 0.96 ± 0.02, 0.94 ± 0.03, 0.91 ± 0.08, 0.91 ± 0.1, 0.97 ± 0.02, 0.95 ± 0.02, 0.95 ± 0.02, respectively. **C)** Locally deformed region near a tear of expanded patterned grid (20 µm squares) with deformation in zoomed region color-coded in magenta as defined by squareness (expansion factor: 8.1). **D)** Pre- and post-expansion images of U2OS cells cultured on GelMap coverslips (grid size: 10 µm, expansion factor: 9.2) stained for tubulin (cyan) and myc-tag (orange). **E)** Schematic of landmark spacing used for correction in F and G. **F)** Quantification of displacement vector length for equidistantly spaced points (1 µm) in the cellular area for uncorrected and pattern-corrected images registered to ground truth tubulin channel. Corresponding deformation fields are color-coded and shown on the right. **G)** Quantification of measurement error (deviation from expected distance assuming isotropic expansion) between pairs of points for a measurement length (mean ± SD). Scale bars: A: 12 mm, B and C: 100 µm, zoom: 40 µm, D and H: 20 µm.

**Supplementary Figure 3:**
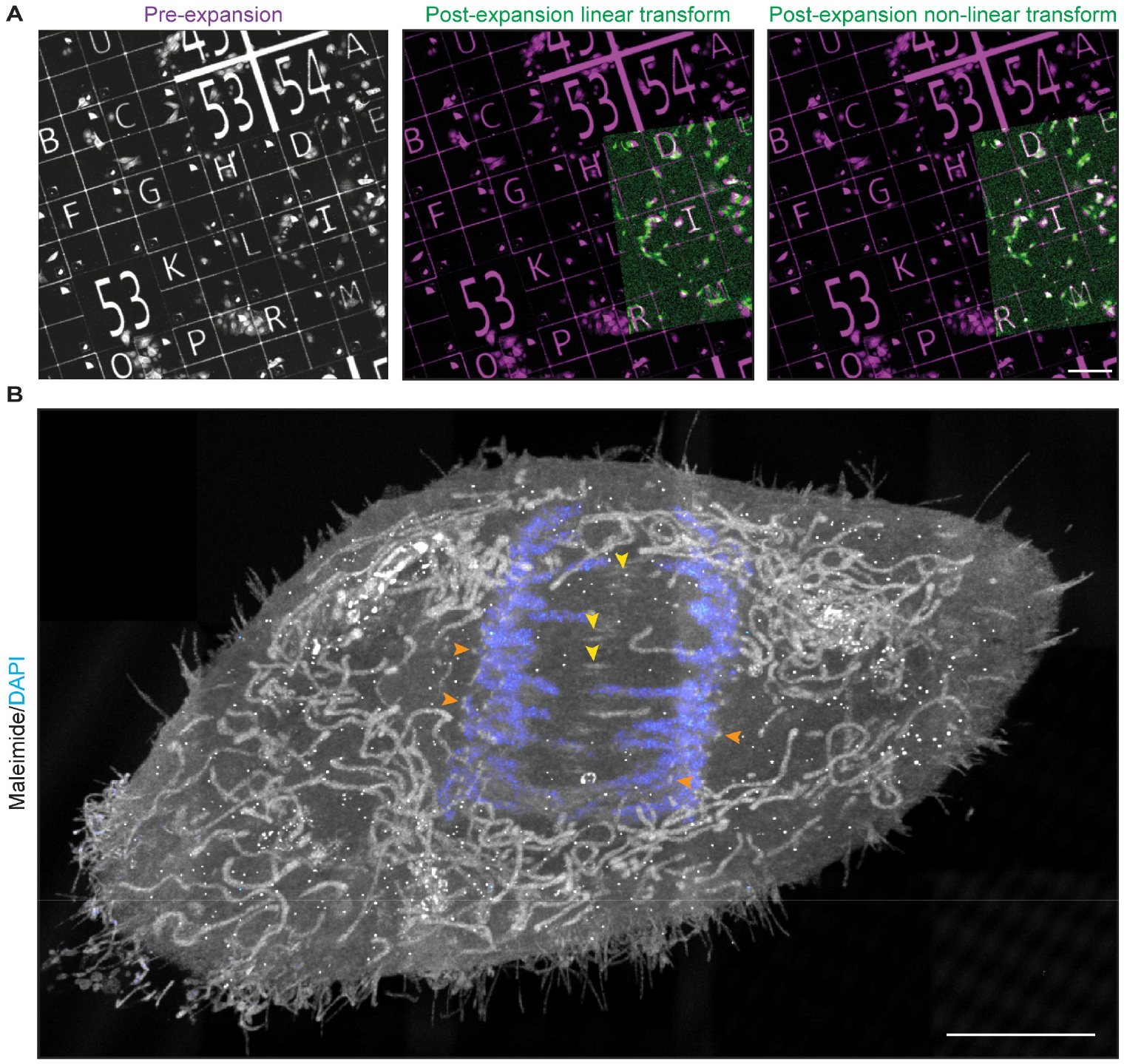
GelMap aids sample navigation and can be used for correction. **A)** Left: pre-expansion image of laminin-594 coated GelMap grid. Middle and right: overlayed linear and non-linear transformed post-expansion images (expansion factor: 10.7). **B)** Additional projection of mitotic cell in Fig. 3D. Scale bars: A: 200 µm B: 5 µm.

**Supplementary Video 1: Correlative Live and Expansion Microscopy using GelMap**

Video corresponding to Fig. 3C-D and Fig. S3B. A U2OS cell stably expressing YFP-H2B and mCherry-H2B and cultured on a GelMap grid was imaged using a spinning disc confocal microscope (30 sec interval) and fixed on the stage during anaphase. The cell of interest was located on the coverslip using the navigation grid, and expanded and imaged using TREx microscopy. Ultrastructural context is provided by maleimide staining, revealing cell morphology, organelle distribution, and spindle components such as centrioles.

## REFERENCES

Bogovic JA, Hanslovsky P, Wong A, Saalfeld S. 2016. Robust Registration of Calcium Images by Learned Contrast Synthesis. 2016 Ieee 13th Int Symposium Biomed Imaging Isbi 1123–1126. doi:10.1109/isbi.2016.7493463

Chang J-B, Chen F, Yoon Y-G, Jung EE, Babcock H, Kang JS, Asano S, Suk H-J, Pak N, Tillberg PW, Wassie AT, Boyden ES. 2017. Iterative expansion microscopy. Nat Methods 14:593 599. doi:10.1038/nmeth.4261

Chen F, Tillberg PW, Boyden ES. 2015. Expansion microscopy. Science 347:543 548. doi:10.1126/science.1260088

Chen G, Deng X. 2018. Cell Synchronization by Double Thymidine Block. Bio-protocol 8. doi:10.21769/bioprotoc.2994

Chen R, Cheng X, Zhang Y, Yang X, Wang Y, Liu X, Zeng S. 2021. Expansion tomography for large volume tissue imaging with nanoscale resolution. Biomed Opt Express 12:5614. doi:10.1364/boe.431696

Damstra HG, Mohar B, Eddison M, Akhmanova A, Kapitein LC, Tillberg PW. 2022. Visualizing cellular and tissue ultrastructure using Ten-fold Robust Expansion Microscopy (TREx). Elife 11:e73775. doi:10.7554/elife.73775

Gambarotto D, Zwettler FU, Guennec ML, Schmidt-Cernohorska M, Fortun D, Borgers S, Heine J, Schloetel J-G, Reuss M, Unser M, Boyden ES, Sauer M, Hamel V, Guichard P. 2019. Imaging cellular ultrastructures using expansion microscopy (U-ExM). Nat Methods 16:71–74. doi:10.1038/s41592-018-0238-1

Gao M, Thielhorn R, Rentsch J, Honigmann A, Ewers H. 2020. Expansion STED microscopy (ExSTED). Methods Cell Biol 161:15–31. doi:10.1016/bs.mcb.2020.06.001

Halpern AR, Alas GCM, Chozinski TJ, Paredez AR, Vaughan JC. 2017. Hybrid Structured Illumination Expansion Microscopy Reveals Microbial Cytoskeleton Organization. Acs Nano 11:12677–12686. doi:10.1021/acsnano.7b07200

Jurriens D, Batenburg V van, Katrukha EA, Kapitein LC. 2020. Methods in Cell Biology. Methods Cell Biol. doi:10.1016/bs.mcb.2020.04.018

Ku T, Swaney J, Park J-Y, Albanese A, Murray E, Cho JH, Park Y-G, Mangena V, Chen J, Chung K. 2016. Multiplexed and scalable super-resolution imaging of three-dimensional protein localization in size-adjustable tissues. Nat Biotechnol 34:973–981. doi:10.1038/nbt.3641

Li H, Warden AR, He J, Shen G, Ding X. 2022. Expansion microscopy with ninefold swelling (NIFS) hydrogel permits cellular ultrastructure imaging on conventional microscope. Sci Adv 8:eabm4006. doi:10.1126/sciadv.abm4006

M’Saad O, Bewersdorf J. 2020. Light microscopy of proteins in their ultrastructural context. Nat Commun 11:3850. doi:10.1038/s41467-020-17523-8

Scheible MB, Tinnefeld P. 2018. Quantifying Expansion Microscopy with DNA Origami Expansion Nanorulers. Biorxiv 265405. doi:10.1101/265405

Schindelin J, Arganda-Carreras I, Frise E, Kaynig V, Longair M, Pietzsch T, Preibisch S, Rueden C, Saalfeld S, Schmid B, Tinevez J-Y, White DJ, Hartenstein V, Eliceiri K, Tomancak P, Cardona A. 2012. Fiji: an open-source platform for biological-image analysis. Nat Methods 9:676–682. doi:10.1038/nmeth.2019

Shaib AH, Chouaib AA, Imani V, Chowdhury R, Georgiev SV, Mougios N, Monga M, Reshetniak S, Mihaylov D, Chen H, Fatehbasharzad P, Crzan D, Saal KA, Trenkwalder C, Mollenhauer B, Outeiro TF, Preobraschenski J, Becherer U, Moser T, Boyden ES, Aricescu ARA, Sauer M, Opazo F, Rizzoli S. 2022. Expansion microscopy at one nanometer resolution. doi:10.1101/2022.08.03.502284

Sim J, Park CE, Cho I, Min K, Eom M, Han S, Jeon H, Cho H-J, Cho E-S, Kumar A, Chong Y, Kang JS, Piatkevich KD, Jung EE, Kang D-S, Kwon S-K, Kim J, Yoon K-J, Lee J-S, Boyden ES, Yoon Y-G, Chang J-B. 2022. Nanoscale resolution imaging of the whole mouse embryos and larval zebrafish using expansion microscopy. Biorxiv 2021.05.18.443629. doi:10.1101/2021.05.18.443629

Théry M. 2010. Micropatterning as a tool to decipher cell morphogenesis and functions. J Cell Sci 123:4201–4213. doi:10.1242/jcs.075150

Tillberg PW, Chen F, Piatkevich KD, Zhao Y, Yu C-C (Jay), English BP, Gao L, Martorell A, Suk H-J, Yoshida F, DeGennaro EM, Roossien DH, Gong G, Seneviratne U, Tannenbaum SR, Desimone R, Cai D, Boyden ES. 2016. Protein-retention expansion microscopy of cells and tissues labeled using standard fluorescent proteins and antibodies. Nat Biotechnol 34:987 992. doi:10.1038/nbt.3625

Truckenbrodt S, Sommer C, Rizzoli SO, Danzl JG. 2019. A practical guide to optimization in X10 expansion microscopy. Nat Protoc 14:832–863. doi:10.1038/s41596-018-0117-3

Valdes PA, Yu C-C (Jay), Aronson J, Zhao Y, Bernstock JD, Bhere D, An B, Viapiano MS, Shah K, Chiocca EA, Boyden ES. 2021. Decrowding Expansion Pathology: Unmasking Previously Invisible Nanostructures and Cells in Intact Human Brain Pathology Specimens. Biorxiv 2021.12.05.471271. doi:10.1101/2021.12.05.471271

Xu H, Tong Z, Ye Q, Sun T, Hong Z, Zhang L, Bortnick A, Cho S, Beuzer P, Axelrod J, Hu Q, Wang M, Evans SM, Murre C, Lu L-F, Sun S, Corbett KD, Cang H. 2019. Molecular organization of mammalian meiotic chromosome axis revealed by expansion STORM microscopy. Proc National Acad Sci 116:18423–18428. doi:10.1073/pnas.1902440116

Yu C-C (Jay), Barry NC, Wassie AT, Sinha A, Bhattacharya A, Asano S, Zhang C, Chen F, Hobert O, Goodman MB, Haspel G, Boyden ES. 2020. Expansion microscopy of C. elegans. Elife 9:e46249. doi:10.7554/elife.46249

Zhao Y, Bucur O, Irshad H, Chen F, Weins A, Stancu AL, Oh E-Y, DiStasio M, Torous V, Glass B, Stillman IE, Schnitt SJ, Beck AH, Boyden ES. 2017. Nanoscale imaging of clinical specimens using pathology-optimized expansion microscopy. Nat Biotechnol 35:757–764. doi:10.1038/nbt.3892

